# A meta reinforcement learning account of behavioral adaptation to volatility in recurrent neural networks

**DOI:** 10.1101/2024.05.09.593363

**Authors:** D. Tuzsus, I. Pappas, J. Peters

## Abstract

Natural environments often exhibit various degrees of volatility, ranging from slowly changing to rapidly changing contingencies. How learners adapt to changing environments is a central issue in both reinforcement learning theory and psychology. For example, learners may adapt to changes in volatility by increasing learning if volatility increases, and reducing it if volatility decreases. Computational neuroscience and neural network modeling work suggests that this adaptive capacity may result from a meta-reinforcement learning process (implemented for example in the prefrontal cortex), where past experience endows the system with the capacity to rapidly adapt to environmental conditions in new situations. Here we provide direct evidence for a meta-reinforcement learning account of adaptation to environmental volatility. Recurrent neural networks (RNNs) were trained on a restless four-armed bandit reinforcement learning problem under three different training regimes (low volatility training only, medium volatility training only, or meta-volatility training across a range of volatilities). We show that, in contrast to RNNs trained in the low volatility regime, RNNs trained under the medium or meta-volatility regimes exhibited a superior adaption to environmental volatility. This was reflected in a better performance, and computational modeling of the network’s behavior revealed a more adaptive adjustment of learning and exploration to varying levels of volatility during test. Results extend the meta-RL account to volatility adaptation and confirm past experience as a crucial factor of adaptive decision-making.

## Introduction

In the field of reinforcement learning (RL) an agent (e.g. an animal, human or artificial agent) learns to optimize actions to maximize reward and minimize punishment, based on feedback signals obtained from the environment (Sutton & Barto, 2018). Rewards increase action probabilities, whereas punishment signals decrease action probabilities.

From a statistical learning perspective, environmental signals have inherent uncertainty if the generating process is based on a latent outcome distribution with some degree of variability (e.g. a Gaussian with mean and variance). At least two types of uncertainty can be distinguished, stochasticity and volatility (Payzan-LeNestour et al., 2013; Payzan-LeNestour & Bossaerts, 2011, 2011; Soltani & Izquierdo, 2019). In stochasticity (or expected uncertainty), the observed variance in outcomes is due to noise inherent in the environment (in the example, due to the variance of the Gaussian). In such a situation, learning should be slow to integrate over all noisy outcomes to approximate the true expected value (Piray & Daw, 2021; Soltani & Izquierdo, 2019). Second, variations in outcomes may be due to changes in contingencies (volatility, e.g., a change in the mean of the reward distribution over time, sometimes also referred to as unexpected uncertainty). Here, unexpected outcomes relate to environmental change, and learning should therefore be accelerated to update value predictions (Behrens et al., 2007; Piray & Daw, 2021; Soltani & Izquierdo, 2019).

To maximize reward in changing environments, an optimal balance between exploration and exploitation is of central importance, as agents face the dilemma of whether to pursue actions that are based on past rewards (exploitation) or to explore new actions for information gain (exploration) (Gershman, 2018, 2019; Schulz & Gershman, 2019; Sutton & Barto, 2018; Wilson et al., 2021). In stable environments (changes in outcomes are noise due to stochasticity), it is expected that agents initially explore all actions and then continuously exploit the most rewarding action. Here learning should be slower and exploitation should be increased. In contrast, in changing environments (changes in outcomes signal contingency changes due to volatility), learning should be faster and exploration should increase.

In standard reinforcement learning (RL) models, learning and exploration are characterized by two parameters, the learning rate and the softmax inverse temperature. The learning rate regulates how rapidly value predictions are updated based on prediction errors (Sutton & Barto, 2018). Both human and animal learners readily adapt learning rates to environmental volatility (Behrens et al., 2008; Cook et al., 2019; Krugel et al., 2009; Massi et al., 2018; McGuire et al., 2014; Nassar et al., 2010; Woo et al., 2023). For example, Behrens et al. (2007) tested participants in a two-armed bandit task consisting of alternating stable and volatile blocks, where the most rewarding action changes often. Human learners reduced learning rates in stable blocks and increased learning rates in volatile blocks, such that they correctly interpreted changes in reward feedback as changes in contingencies (volatility), and consequently updated value predictions faster in volatile vs. stable blocks. The inverse temperature (β or softmax slope) regulates the balance between exploitation and exploration (Brown et al., 2022; Harada, 2020; Sutton & Barto, 2018; Wilson et al., 2014). Higher values of β reflect exploitation, whereas random exploration increases as β decreases (for β=0, choices are completely random). Adaptive behavior under increasing volatility would therefore entail increasing learning rates, and decreasing β (increasing exploration).

Alterations in learning under uncertainty, due to volatility and stochasticity, are linked to several psychiatric disorders including anxiety (Aylward et al., 2019; Huang et al., 2017), depression (Pulcu & Browning, 2019), obsessive-compulsive disorder (Apergis-Schoute & Ip, 2020; Moreira et al., 2020), substance use disorder (Zuhlsdorff, 2022) and gambling disorder (Addicott et al., 2015; Jara-Rizzo et al., 2020; Wiehler et al., 2021). For example, anxiety (clinical and non-clinical) is thought to be related to a maladaptive response to stochasticity, such that stochastic variations of reward outcomes are misinterpreted as meaningful environmental changes (Aberg et al., 2022; Aylward et al., 2019; Browning et al., 2015; Huang et al., 2017; Pulcu & Browning, 2019). Such effects can be captured by computational models that estimate stochasticity and volatility based on reward feedback (Piray & Daw, 2021). Here, lesioning the model’s stochasticity module reproduced maladaptive learning rate adaptations as observed in anxiety, reinforcing the link between uncertainty misestimation and psychiatric disorders (Pulcu & Browning, 2019).

Past experience of volatility may be a primary driver of adaptive behavior (Egner, 2023; Nassar & Troiani, 2021; Yu et al., 2021). Here, previous work has often focused on cognitive control, i.e. the ability to dynamically adjust the balance between cognitive stability and flexibility to environmental demands. In RL terms, cognitive stability is conceptually related to lower learning rates and higher inverse temperature in stable environments, whereas cognitive flexibility is conceptually related to higher learning rates and lower inverse temperature observed under high volatility. Such adaptive behavior may result from a reinforcement learning process (Egner 2023) that adjusts control resources based on a recent history of task demands. Reinforcing this idea, human learners can indeed learn environment-specific learning rates (Simoens et al., 2023) and past experience of volatility modulates this ability (Wen et al., 2023). Past experience of volatility may also have more general benefits for reinforcement learning under uncertainty (Hohl & Dolcos, 2024). This idea resonates well with primates showing enhanced encoding of reward signals in prefrontal and orbitofrontal cortex for volatile vs. stable conditions (Massi et al., 2018) and human learners adapting their learning rate if transitioning from volatile to stable but not from stable to volatile environments (Xu et al., 2021).

The adaptation of cognitive control to novel task demands is also referred to as meta-control (Egner, 2023). One computational model for this meta control is meta-RL (Wang et al., 2018). In the meta-RL framework, agents (e.g. neural network models) are trained on samples from a given task distribution to learn a representation of the task that allows them to generalize to novel task instantiations. Crucially, in this framework, neural network weights are updated only during training. During test, weights are held fixed, and any adaptive behavior arising during test is attributable to the computational mechanism embedded in the network. Some accounts suggest that this meta-RL capacity might be implemented in prefrontal circuits (Wang et al., 2018). RNNs trained using a meta-RL approach showed an adaptation of learning rates to environmental volatility (Wang et al. 2018), similar to human learners (Behrens et al., 2007). This flexibility arose without retraining (e.g., when weights and biases were held fixed during test). However, the role of past experience as well as the adaptation to varying levels of volatility during test was not examined.

Here we directly tested the hypothesis that volatility adaptation arises from a meta-RL process, where previous experience in terms of volatility of past tasks is a critical driver. To test this hypothesis, we examined the effect of exposure to various levels of volatility during training on volatility adaptation during test in RNNs. Specifically, we applied three training regimes. One set of networks was trained only on low volatility environments. A second set of networks was trained only on medium volatility environments. A final set of networks was trained using a meta volatility scheme, were training involved episodes covering a large range of volatilities. Based on the considerations outlined above, adaptation to volatility was quantified by examining the capacity of networks to flexibly adjust learning rates and inverse temperature parameters to changing volatility levels during test.

## Methods

### Network Architecture

For each training regime (see below) n=30 RNN models based on LSTM units endowed with computational noise (Findling & Wyart, 2020; Tuzsus et al., 2024) where trained. Standard LSTM dynamics (Hochreiter & Schmidhuber, 1997) were characterized by following equations:

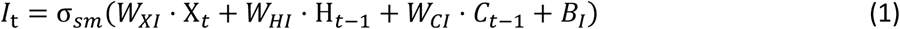

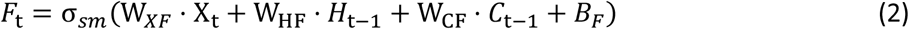

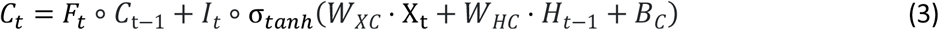

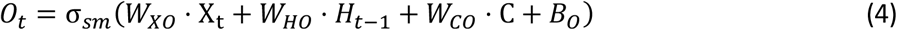

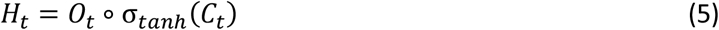

*X* represents the input, which is a one-hot encoding of the previous action and the previous reward. *I* denotes the input gate, *F* corresponds to the forget gate (alternatively known as the maintenance gate), C represents the cell state, *O* is the output gate, *H* is the hidden state, and *t* indexes trials. *σ*_*sm*_ and *σ*_*tanh*_ refer to the softmax or hyperbolic tangent activation functions, respectively. The network’s trainable parameters consist of weight matrices *W* and bias vectors *B*, with subscripts indicating the associated gates or states.

Computational noise following Weber’s law of intensity sensation accounts for adaptive behavior and neural signals in human learners exposed to volatile RL tasks (Findling et al., 2019; Findling & Wyart, 2021). Importantly, such computational noise was also shown to improve adaptation of RNN behavior to adverse RL environments (e.g., tasks with increased volatility)(Findling & Wyart, 2020), which allows them to perform as good as human learners in the exact same volatile bandit task (Tuzsus et al., 2024).

Therefore, following Findling & Wyart (2020) we introduced noise (*Weber noise*) at the level of the hidden state *H* of an LSTM unit *k* at trial *t*:

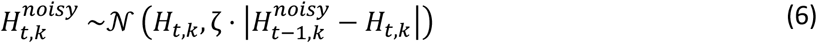

### Restless Bandit Task

Agents were trained on a 4-armed restless bandit task (Daw et al., 2006) for 50.000 episodes, where each episode consisted of 300 trials. In the restless bandit task, the reward of the *i*th action in trial *t* is the reward of the same action from the preceding trial t-1, plus noise:

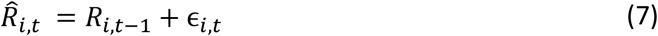

Noise ϵ was drawn from a gaussian distribution with mean 0 and standard deviation *σ*:

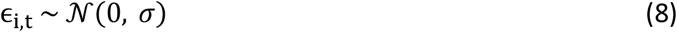

We implemented reflecting boundaries to ensure that reward was within [0,1]:

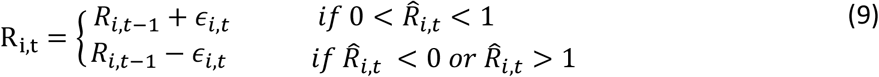

### Training regimes

To test the hypothesis that volatility adaptation arises from a meta-RL process, we tested the effect of exposure to various levels of volatility during training on volatility adaptation during test in RNNs. We therefore applied three training regimes (n=30 networks each). One set of networks was trained only on low volatility environments, here *σ* (Eq. 8) was set 0.05. A second set of networks was trained only on medium volatility environments, here *σ* was set to 0.1. A final set of networks was trained using a meta volatility scheme, where in each training episode *σ* was drawn from a uniform distribution with a lower boundary of 0.02 and an upper boundary of 0.2.

### Test

Following the standard meta-RL approach, after training, network weights were fixed, and the network performance under various levels of volatility was examined. For all network instances, testing used the same set of ten runs for each volatility level (15 values of *σ* ranging from 0.02 to 0.3 in 0.02 steps).

### Behavioral Analysis

Model-agnostic behavioral measures included the percentage of optimal choices (i.e., selecting the most rewarding action on a given trial) and the percentage of switches (i.e., selecting a different action than the action of the previous trial).

To investigate behavioral results across different test volatilities, we aggregated the percentage of optimal choices and the percentage of switch trials for each network instance and test volatility training regime. Subsequently, mean values of aggregated data were tested for main effects of training volatility using Bayesian ANOVAs and post-hoc independent Bayesian t-tests as implemented in JASP (JASP Team, 2024). To quantify evidence for the alternative hypothesis (e.g., difference in means of training volatility regimes) relative to the null hypothesis we used Bayes Factors (BF, Beard et al., 2016). Here we used established BF cut-offs were the degree of evidence for the alternative hypothesis is characterized by the following values: BF > 100: decisive evidence; 100-30: very strong evidence; 30-10: strong evidence, 10-3: moderate evidence; 3-1: anecdotal evidence. For readability we used subscripts BF_10_ for evidence regarding the alternative hypothesis and BF_01_, the inverse Bayes Factor, for evidence in favour of the null hypothesis.

Although analyses on the aggregate level characterize main effects of training regime, this approach fails to provide information on how networks adapt to changes in test volatilities. To formally characterize the relationship between test volatility and performance, we fitted behavioral measures with an exponential decay function over test volatilities via an exponential decay function fitted to the median values per test volatility level, separately for the three different training regimes:

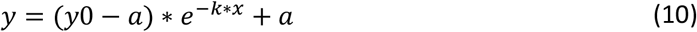

Here, *y* is the model-agnostic behavioral measure (% optimal or % switch) for each level of test volatility *x* (i.e., the standard deviation of the noise distribution *σ*, see Equation 8), *y*0 is the intercept (e.g., the value of *y* for a test volatility of 0.02), *a* is the asymptote of the function, and *k* is the decay rate.

Function parameters were estimated via Hamiltonian Monte Carlo sampling (2 chains, 30.000 samples, warmup = 15000) using Stan, and the Rstan package in R (Version 4.1.1). Convergence was assumed if 1 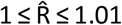 (Gelman & Rubin, 1992). Effects were considered reliable if the 95% highest density interval of the difference in posterior distributions between conditions did not include zero. To further characterize effects, we additionally report directional Bayes Factors (dBFs) (Marsman & Wagenmakers, 2017) to quantify the degree of evidence for differences in parameter values for *y0, a* and *k* between the three training regimes. dBFs were calculated as the ratio of the integral of the difference distribution of the respective parameters between training volatility regimes from −∞ to 0 (negative effect integral) versus the integral from 0 to ∞ (positive effect integral). To aid interpretability we denoted dBF_pos_ if the positive effect integral was in the numerator and dBF_neg_ if the negative effect integral was in the numerator. A dBF_pos_ of 5 thus indicates that a positive directional effect was five times more likely than a negative directional effect, in contrast dBF_neg_ of the same value would indicate that a negative effect was five times more likely than a positive directional effect.

### Computational Modelling

We also modelled the data using a standard Q-learning RL model, where agents update the expected value 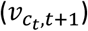 on trial t+1 of the chosen action based on the prediction error *δ*_*t*_ experienced on trial *t*:

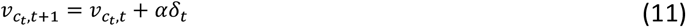

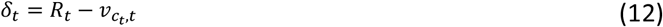

The learning rate 0 ≤ α ≤ 1 regulates the proportion of the prediction error utilized for updating. *R*_*t*_ represents the reward obtained on trial *t*. Unchosen action values remain unchanged. Values were initialized to *v*_*c*,0_ = 0.5.

Action values *v* were transformed into choice probabilities via a standard softmax function:

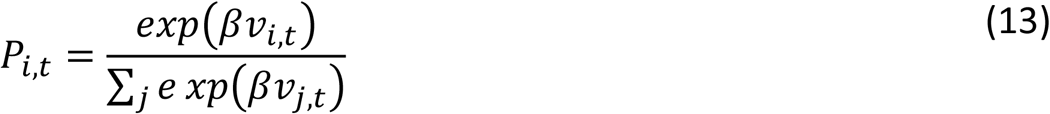

*p*_*i*,*t*_ is the probability of selecting action *i* on trial *t*, and β is the inverse temperature parameter that governs exploration. Lower values of β correspond to higher levels of choice stochasticity (i.e. higher levels of random exploration).

Models were fit to each agent’s behavioral data using Maximum-Likelihood estimation in R (Version 4.1.1) with the optim function. Analysis of fitted model parameters mirrored behavioral data analysis. First, we aggregated learning rates and inverse temperature parameters for each network instance and training regime and examined average values of aggregated data for main effects of training regime with Bayesian ANOVAS and follow-up Bayesian t-tests. Second, we analysed parameter estimates as a function of test volatility by fitting an exponential decay function (See Equation 10) to median parameter estimates per test volatility level, separately for the three different training regimes.

## Results

### Model-agnostic behavioral results

In a first step, we compared behavioral performance between training regimes pooling across levels of test volatility. This revealed decisive evidence for an effect of training volatility of the percentage of optimal choices (Bayesian ANOVA: BF_10_ > 100, Figure 2A). Post Hoc Bayesian t-tests revealed decisive evidence for superior performance of both medium (BF_10_ > 100) and meta volatility trained RNNs (BF_10_ > 100) compared to networks trained on low volatility. In contrast, there was strong evidence against a performance meta and medium volatility trained RNNs (BF_01_ = 10.213). RNNs trained with the medium and meta-volatility regimes therefore exhibited superior performance overall. We next examined switching behavior, i.e. the proportion of trials on which a different option from the previous trial was chosen (a proxy for exploration). This again revealed decisive evidence for an effect of training volatility (BF_10_ > 100, Figure 2B). Post Hoc Bayesian t-tests yielded decisive evidence for reduced switching behavior in networks trained with low volatility compared to medium (BF_10_ > 100) or meta volatility training schemes (BF_10_ > 100). There was very strong evidence that meta volatility trained RNNs showed more switching behavior than those trained with medium volatility (BF_10_ = 40.664). RNNs trained using the meta-volatility regime showed highest level of exploration in terms of switches across all test volatilities, followed by RNNs trained with medium volatility. RNNs trained with low volatility showed the lowest levels of exploration.

**Figure 1:**
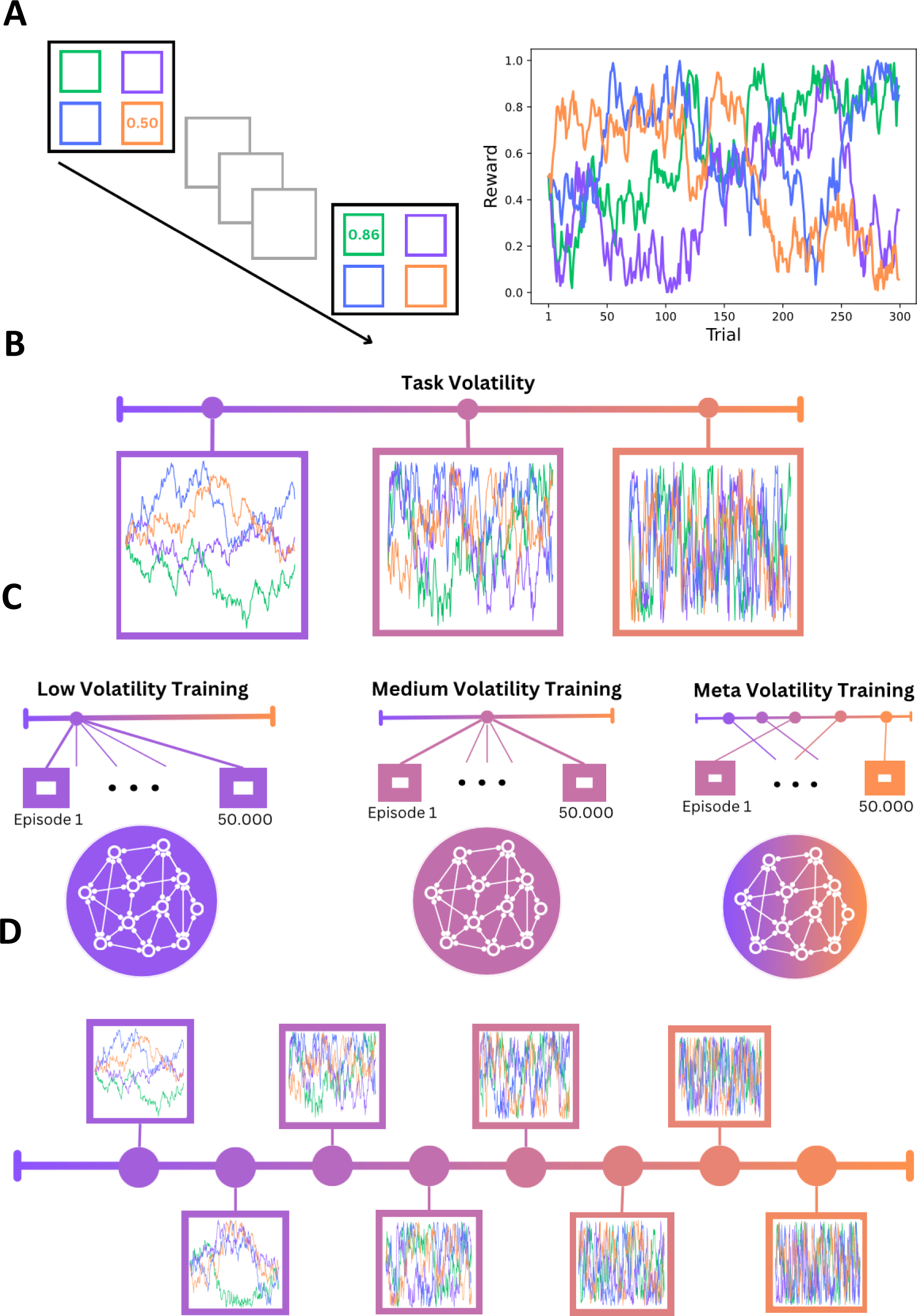
A) Restless Bandit Task: In each trial an agent selects one out of four actions with rewards varying between trials according to independent random walk processes. B) Task Volatility is characterized by the speed of change in random walks, which can range from very stable environments (purple) to very volatile environments (orange). C) Artificial Recurrent Neural Networks (RNNs) were trained either in low volatility, medium volatility or on a range of volatilities (Meta Volatility). D) After training, all RNNs were exposed to novel task instances ranging from low to high volatility.

**Figure 2:**
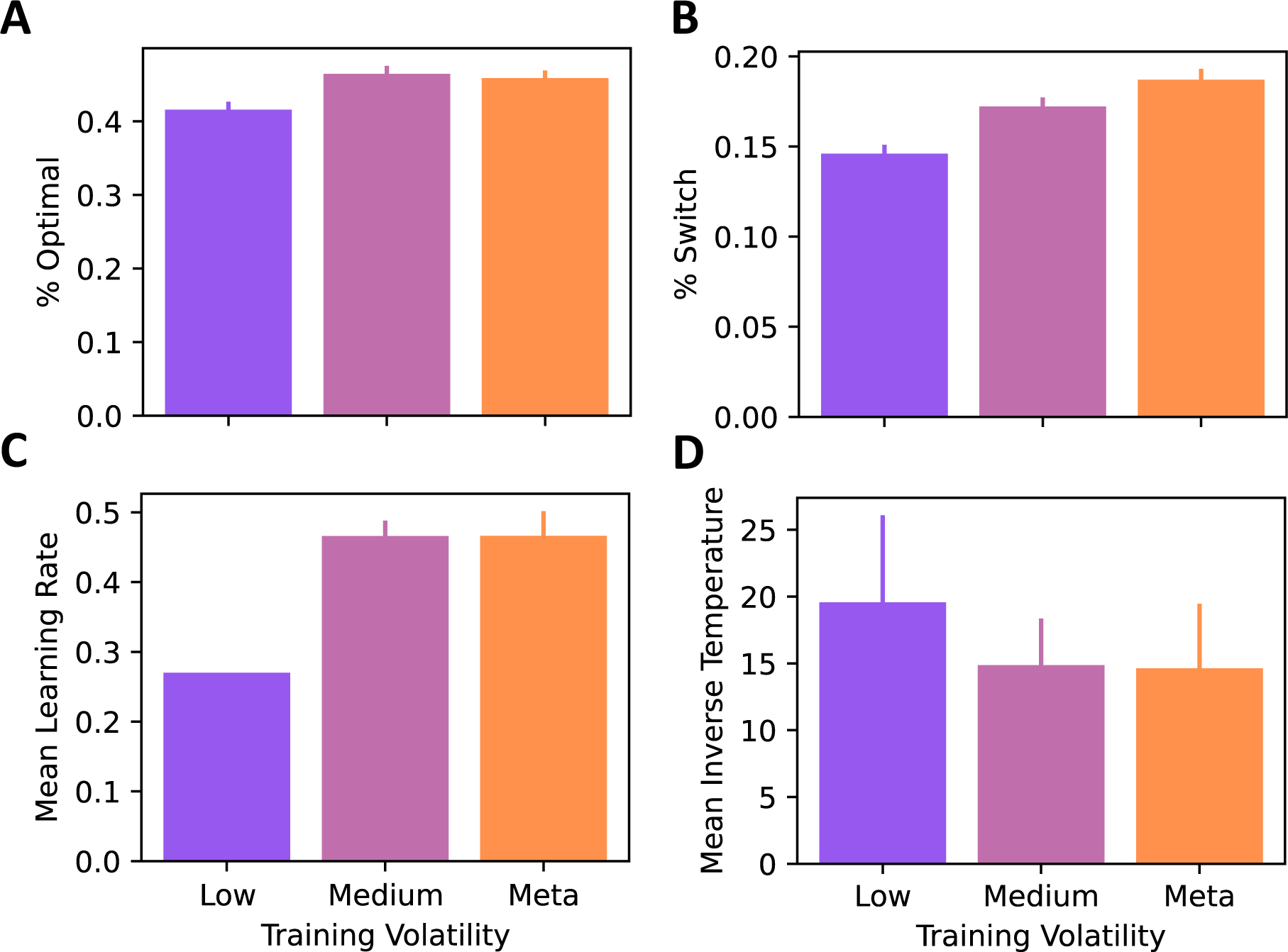
Analysis of Variance of aggregated data. Panels (A: % Optimal Choices, B: % Switch, C: Mean learning rate, D: Mean inverse temperature) correspond to the mean value across test volatility for each training volatility level. Error bars denote 95% confidence intervals. Error bar in C is very low.

Although the aggregate analysis above revealed main effects of volatility training regimes across test volatility levels, it fails to provide more fine-grained information on volatility adaptation. For example, with increasing volatility, one would predict that performance gradually decreases, and switches increase, as due to increasing exploration demands. To investigate differences in adaptation to test volatility between training regimes, we fitted an exponential decay function to the dependent variables (performance and switch rate, see Figure 3A and B), yielding estimates of offset (y0), decay rate (k) and asymptote (a) (see Equation 10). Medians and 95% highest posterior density intervals (HDIs) of the parameter difference distributions are listed in Supplemental Table 1. Differences were considered reliable if zero fell outside the 95%HDI of the difference distribution.

**Figure 3:**
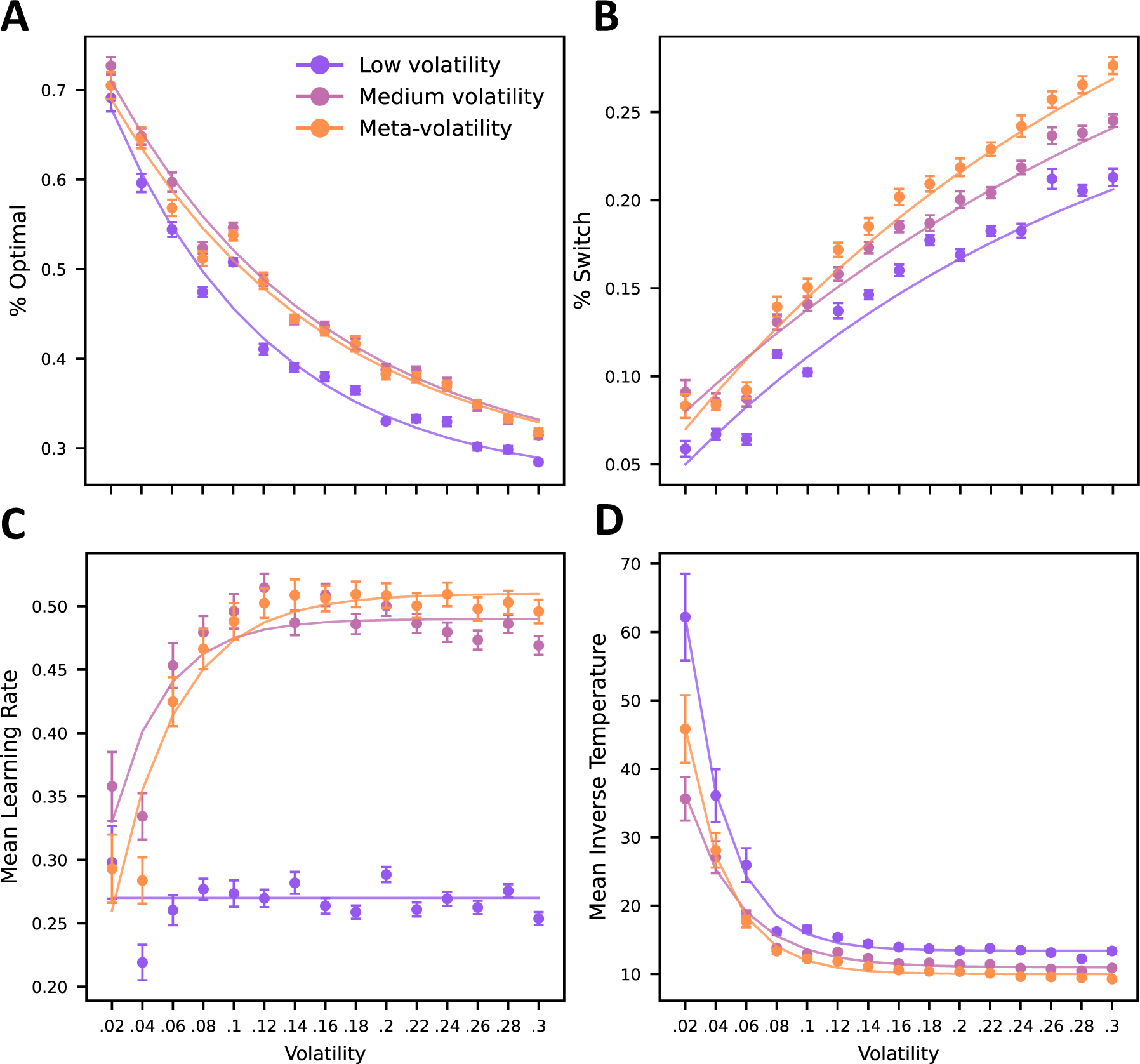
Behavioral (A: % Optimal Choices, B: % Switch) and cognitive adaptation regarding delta-rule parameters (C: Mean learning rate, D: Mean inverse temperature) to test volatility (x-axis) for agents trained with different volatility schedules (see legend). Points denote means and error bars denote 95% confidence intervals. Lines denote optimal fits of an exponential decay function to the means.

Regarding performance (Figure 3A), no credible parameter differences between training volatility regimes emerged, although both y0 and a were numerically reduced in the low volatility scheme (see Supplemental Table 1). Regarding the switch rate, a higher intercept (*y0*) was observed for medium (95% HDI: [0.004 - 0.051], dBF_pos_ = 78.841) and and meta-volatility regimes (95% HDI: [0.001 - 0.045], dBF_pos_ = 28.017), compared to the low volatility training regime, reflecting lower baseline exploratory behavior in networks trained under low volatility.

In summary, compared to RNNs trained with the low volatility regime, RNNs trained with meta or medium volatility exhibited better performance, and increased exploration behavior in terms of switch rate.

### Modeling Results

To directly test our computational hypotheses regarding learning rate and softmax inverse temperature, RNN behavioral data were fitted with a simple delta rule RL model (see Equations 11-13). As in the model-agnostic behavioral analyses (see previous section), we first examined overall effects by aggregating parameters (learning rates, Figure 2C, and inverse temperatures see Figure 2D) across test volatilities. For mean learning rates there was decisive evidence for a main effect of training volatility (Bayesian ANOVA: BF_10_ > 100). Post Hoc Bayesian t-tests revealed decisive evidence for lower learning rates in low versus medium (BF_10_ > 100) and low vs. meta volatility trained RNNs (BF_10_ > 100). There was only anecdotal evidence for a difference between medium and meta volatility trained networks. This strongly suggests that, in stark contrast to RNNs trained in the low volatility regime, RNNs trained in the medium volatility or meta-volatility regime flexibly adjusted learning rates to changing test volatilities. Aggregated inverse temperature values showed only anecdotal evidence for a main effect of training volatility (BF?), likely due to the high variance in inverse temperature estimates (see error bars in Figure 2C).

As in the behavioral analysis above, to gain fine-grained information of adaptation mechanisms to test volatility, model parameters were fitted with an exponential decay function (Equation 10) across levels of test volatility. The medians and highest posterior density intervals of the difference distributions of function parameters are listed in Supplemental Table 1. RNNs trained in the medium and meta volatility regimes showed a higher learning rate asymptote than RNN trained using low volatility (medium vs. low: 95% HDI: [0.112, 0.322], dBF_pos_ = 46.48, meta vs. low: 95% HDI: [0.130, 0.340], dBF_pos_ = 50.05), reflecting the fact that low volatility trained networks failed to increase learning rates in high volatility environments.

RNNs trained in medium or meta volatility regimes also showed increased baseline exploration at lower test volatilities, which was reflected by lower intercept (y0) compared to low volatility trained networks (see Figure 3C, medium: 95% HDI: [−28.723, −23.249], dBF_neg_ = Inf; meta: 95% HDI: [−18.521, −13.694], dBF_neg_ = Inf). This mirrors the higher baseline switch rates observed in the behavioral analysis above (Figure 3B). Furthermore, these RNNs also showed an improved ability to dynamically adapt exploration to test volatility, which was reflected in a lower inverse temperature asymptote (meta: 95% HDI: [−4.220, −2.598], dBF_neg_ = 49048.67, medium: 95% HDI: [−3.402, −1.490], dBF_neg_ = 7235.039) and lower decay rate in the medium volatility regime (95% HDI: [−0.283, −0.057], dBF_neg_ = 268.107). RNNs trained with medium or meta-volatility showed increased baseline exploration and a superior ability to increase exploration depending on environmental demands. Regarding exploration, several differences between medium- and meta-volatility training schemes emerged. Meta-volatility trained RNNs exhibited less exploration in more stable environments (higher intercept *y0*: 95% HDI: [7.518, 11.875], dBF_pos_ = Inf), but more exploration in higher volatility environments (lower asymptote *a*: 95% HDI: [7.518, 11.875], dBF_neg_ = 107.328). They also adjusted exploration more rapidly to increasing test volatilities (higher decay rate *k:* 95% HDI: [0.045, 0.255], dBF_pos_ = 150.137), leading to an overall more flexible adaptation of exploration behavior to test volatility levels in networks that experienced various environmental dynamics during training.

In summary, in contrast to networks trained with low volatility training regime, networks trained with the medium or meta volatility regimes dynamically adjusted learning rates to environmental demands. A complementary pattern was observed for exploration (inverse temperature): RNNs trained with low volatility showed reduced exploration. In contrast, meta volatility networks explored more (lower intercept *y0* and asymptote *a*) and more flexibly adjusted exploration to environmental demands than RNNs trained in the medium or low volatility regimes.

## Discussion

Human and non-human learners readily adapt decision-making to environmental dynamics (e.g., volatility) (Behrens et al., 2007, 2008; Cook et al., 2019; Krugel et al., 2009; Massi et al., 2018; McGuire et al., 2014; Nassar et al., 2010; Woo et al., 2023), and there is some evidence for similar adaptation mechanisms in recurrent neural networks (RNNs) (Wang et al., 2018). Theoretical and empirical work suggests that past experience of volatility could be a critical driver for rapid adaptation to new task demands (Cook et al., 2019; Egner, 2023; Nassar & Troiani, 2021; Wen et al., 2023; Yu et al., 2021). Here we directly tested the hypothesis that volatility adaptation arises from such a meta-RL process by training RNNs as standalone artificial agents on four-armed restless bandit tasks (Chakroun et al., 2020; Daw et al., 2006; Speekenbrink & Konstantinidis, 2015; Wiehler et al., 2021). Expanding upon previous work (Wang et al 2018), we directly examined the role of previous experience by applying three training regimes. One set of RNNS was trained only on low volatility environments. A second set of RNNs was trained only on medium volatility environments, and a final set was trained using a meta volatility scheme, were training involved episodes covering a large range of volatilities. Furthermore, adaptation to environmental volatility during test was examined by testing networks over a broad range of volatility levels, allowing us to comprehensively examine the networks’ adaptation behavior.

In a first step, behavioral adaptation was examined in terms of model-agnostic measures, i.e. performance (% optimal choices) and switch rate. This confirmed the central role of training experience: aggregated over all test volatility levels (Figure 2A), RNNs trained with meta or medium volatility showed superior performance compared to networks trained using the low volatility regime, in particular under high volatility test conditions (see Figure 3A). Aggregated over all test volatility levels, RNNs trained with meta or medium volatility regimes also explored more (see Figure 2B), which was reflected in a higher switch rate asymptote and intercept compared to RNNs trained on low volatility (see Figure 3B).

Improved performance of RNNs trained with medium and meta-volatility resonate well with animal and human studies. For example, studies in non-human primates show that volatility facilitates value-based learning (Massi et al. 2018), such that animals show a more robust encoding of reward signals in PFC and OFC under volatile compared to stable task conditions. Relatedly, Xu et al. (2021) showed that in human learners learning adjustments are more pronounced when transitioning from volatile to stable, but not from stable to volatile environments. In Wen et al. (2023), participants were either trained under high or low volatility conditions in the Wisconsin card sorting task, with frequent or infrequent rule changes, respectively. Resonating with our findings in RNNs, participants trained in low volatility environments were more likely to stick to past task rules and were slower to adapt to task rule changes than participants trained under high volatility.

The second level of analysis was adaptation in terms of learning rate and inverse temperature parameters from fitting a delta-rule RL model. Theoretical and empirical work suggests that learning rates should increase under conditions of high volatility, to allow for a rapid adjustment to changes in the environment (Behrens et al., 2007; Piray & Daw, 2021; Soltani & Izquierdo, 2019). Aggregated across all task volatility levels, there was decisive evidence for reduced learning rates in RNNs trained on low volatility compared to RNNs trained with medium or meta volatility (see Figure 2C). This was due to these networks’ ability to flexibly adjust learning rates to test volatility. In contrast, learning rates during test were highly rigid for low volatility trained RNNs (see Figure 2C). Similar adaptations to environmental volatility via a modulation of learning rates were shown in several human and non-human studies (Behrens et al., 2007, 2008; Cook et al., 2019; Krugel et al., 2009; Massi et al., 2018; McGuire et al., 2014; Nassar et al., 2010; Soltani et al., 2006; Woo et al., 2023) and in RNNs (Wang et al., 2018). The present results show that this adaptive capacity crucially depends on previous experience. A potential neuronal mechanism underlying the adaptation of learning rates could be implemented by neuronal information integration over different temporal time scales (e.g., learning rates) (Danskin et al., 2023). For example, one animal study has shown that the retrosplenial cortex acts as a reservoir of exponential history integrators, where neurons encode history information with different integration windows (Danskin et al., 2023). Past experiences of different volatility levels may be required to establish these different integrators populations, which could then be dynamically recruited according to the current task demands.

Maladaptive adaptation to volatility is a core feature of several psychiatric disorders, including anxiety (Aylward et al., 2019; Huang et al., 2017), depression (Pulcu & Browning, 2019), obsessive-compulsive disorder (Apergis-Schoute & Ip, 2020; Moreira et al., 2020), substance use disorder (Zuhlsdorff, 2022) and gambling disorder (Addicott et al., 2015; Jara-Rizzo et al., 2020; Wiehler et al., 2021). A common observation in affective disorders such as anxiety and depression is that learning rates are not adapted to the true volatility of the environment (Brown et al., 2023; Pulcu & Browning, 2019). Cognitive control required for adaptation could be generalized from previous experiences of volatility (Hohl et al 2023), resonating with our neural network modeling results. Whether volatility training increases learning rate adaptability in psychiatric disorders is a question for future research. In biological systems, adaptation to environmental volatility might be linked to catecholamine functioning, which is distorted in many psychiatric disorders. For example, exploration behavior depends on dopaminergic neurotransmission, and enhancing catecholamine availability improves learning rate adjustments to task conditions (Cook et al., 2019). Volatility training may mimic this effect and rehabilitate the ability to adapt learning rates in affective disorders such as anxiety and depression.

We quantified exploration via the inverse temperature parameter (Brown et al., 2022; Harada, 2020; Wilson et al., 2014), reflecting random exploration (Sutton & Barto 2018). Across all training regimes, networks adapted to increasing volatility by decreasing inverse temperatures (increasing exploration, see Figure 3D). However, RNNs trained on meta or medium volatility showed strong evidence increased exploration (the exploration intercept and asymptote were lower) compared to RNNs trained in the low volatility regime. Note that the main effect of inverse temperature aggregated over all test volatility levels (see Figure 2D) was inconclusive, likely due to high variance of estimates. Human learners exhibit lower inverse temperature in volatile versus stable environments (Wen et al., 2023; Zhou et al., 2024), resonating with the observations in RNNs. However, there are trade-offs between learning rates and inverse temperature parameters in RL models, which should be kept in mind when interpreting these results (Ballard & McClure, 2019). Prior exposure to volatility endows a network with increasing baseline levels of exploration, which improves performance across a wide range of volatilities. Taken together, modeling results show that networks trained with medium or meta-volatility show a superior adaptation to environmental dynamics, such that learning is slower and exploration is reduced in stable environments, whereas learning is faster and exploration increases in more volatile environments.

Several limitations need to be addressed. First, we focused on the effects of training regime on behavioral adaptation to volatility in one specific neural network architecture (a 48-unit LSTM network trained with the A2C algorithm (Mnih et al., 2016) and *Weber noise* (Findling & Wyart, 2020)), based on our previous results that this architecture exhibits human-level performance on four-armed restless bandit problems (Tuzsus et al., 2024). Previous work on meta-RL applied similar RNN architectures (Binz & Schulz, 2022; Blanco-Pozo et al., 2024; Findling & Wyart, 2020; Hattori et al., 2023; Molano-Mazón et al., 2023; Wang et al., 2017, 2018), but generally, conclusions are limited to the architectures examined. Also, architectures, with mechanisms such as attention (e.g. transformer-based architectures) could be promising for future work (Chen et al., 2021; Parisotto et al., 2020; Schubert et al., 2024; Upadhyay et al., 2019). Second, we focused on volatility (unexpected uncertainty), whereas stochasticity (expected uncertainty), was set to zero (there was no observation noise). Future studies could either focus on stochasticity and/or stochasticity-volatility interactions in training and test, as both affect decision making (Bland & Schaefer, 2012; Payzan-LeNestour & Bossaerts, 2011; Piray & Daw, 2021; Soltani & Izquierdo, 2019) and may have contrasting effects on e.g. learning rates (Piray & Daw, 2021). Stochasticity estimates may also serve as a baseline for the detection of volatility (Soltani & Izquierdo, 2019), which could be tested directly in RNNs via different training regimes (Soltani & Izquierdo, 2019). Third, we focused on a restless bandit tasks used in many previous human and non-human studies (Chakroun et al., 2020; Daw et al., 2006; Ebitz et al., 2018; Speekenbrink & Konstantinidis, 2015; Wiehler et al., 2021) as well as modeling RNN studies (Findling & Wyart, 2020; Tuzsus et al., 2024; Wang et al., 2017). Future work is required to extend these findings to other tasks used to study learning rate adaptation (Behrens et al., 2007, 2008; Browning et al., 2015; Cook et al., 2019; Gagne et al., 2020) (Wang et al., 2018). Finally, computational modeling was restricted to an RL model, that disregarded effects of higher-order choice perseveration (Lau & Glimcher, 2008; Miller et al., 2019; Tuzsus et al., 2024) and more complex directed (i.e. uncertainty-guided) exploration mechanisms (Chakroun et al., 2020; Wiehler et al., 2021; Wilson et al., 2014). However, at least in the network architecture examined here, random exploration appears to be prominent, and directed exploration plays a lesser, if any, role (Tuzsus et al., 2024). Nonetheless, future studies might benefit from examining a more extended computational model space.

Taken together, here we provide evidence for a meta-RL account of behavioral adaptation to changing environmental demands in RNNs. A comparison of three training regimes involving exposure to low volatility, medium volatility or a range of volatilities (meta-volatility) revealed that networks exposed to volatility during training showed superior performance across a range of test volatilities. In line with a meta-RL account, this dynamic adaptation occurred without further weight changes during test (Wang et al., 2018). Computational modelling confirmed that this effect was attributable to a dynamic adjustment of learning rates and exploration behavior to environmental demands, resonating with previous results from human and animal research. Overall, this suggests that past experience is a prerequisite for adaptive decision making in changing environments.

## Acknowledgements

This work was funded by Deutsche Forschungsgemeinschaft (PE1627/8-1, Code 496990750 to J.P.).

## Supplement

**Supplemental Table 1:** Medians and 95% HDIs of the posterior difference distributions of low, medium and meta volatility trained RNNs for the parameters of the exponential decay function.

| DV | Parameter | Meta – Low | Directional BF | Medium - Low | Directional BF | Meta – Medium | Directional BF |
| --- | --- | --- | --- | --- | --- | --- | --- |
| % Optimal | $y_0$ | 0.016 (-0.028, 0.060) | $\text{dBF}_{\text{pos}} = 3.225$ | 0.035 (-0.008, 0.079) | $\text{dBF}_{\text{pos}} = 16.122$ | -0.019 (-0.060, 0.020) | $\text{dBF}_{\text{neg}} = 4.873$ |
| | $a$ | 0.010 (-0.085, 0.095) | $\text{dBF}_{\text{pos}} = 1.525$ | 0.007 (-0.084, 0.089) | $\text{dBF}_{\text{pos}} = 1.334$ | 0.003 (-0.098, 0.101) | $\text{dBF}_{\text{pos}} = 1.117$ |
| | $k$ | -0.043 (-0.123, 0.035) | $\text{dBF}_{\text{neg}} = 6.384$ | -0.043 (-0.118, 0.035) | $\text{dBF}_{\text{neg}} = 6.769$ | -0.001 (-0.071, 0.069) | $\text{dBF}_{\text{neg}} = 1.046$ |
| % Switch | $y_0$ | **0.021 (0.001 to 0.045)** | **$\text{dBF}_{\text{pos}} = 28.017$** | **0.027 (0.004 to 0.051)** | **$\text{dBF}_{\text{pos}} = 78.841$** | -0.006 (-0.026 to 0.016) | $\text{dBF}_{\text{neg}} = 2.499$ |
| | $a$ | 0.109 (-0.334 to 0.507) | $\text{dBF}_{\text{pos}} = 4.354$ | 0.091 (-0.297 to 0.619) | $\text{dBF}_{\text{pos}} = 3.221$ | 0.016 (-0.473 to 0.380) | $\text{dBF}_{\text{pos}} = 1.205$ |
| | $k$ | -0.011 (-0.089 to 0.065) | $\text{dBF}_{\text{neg}} = 1.563$ | -0.021 (-0.100 to 0.062) | $\text{dBF}_{\text{neg}} = 2.217$ | 0.001 (-0.057 to 0.071) | $\text{dBF}_{\text{pos}} = 1.602$ |
| Learning Rate | $y_0$ | -0.014 (-0.077, 0.049) | $\text{dBF}_{\text{neg}} = 2.079$ | 0.057 (-0.012, 0.120) | $\text{dBF}_{\text{pos}} = 18.716$ | -0.071 (-0.143, 0.004) | $\text{dBF}_{\text{neg}} = 32.293$ |
| | $a$ | **0.244 (0.130, 0.340)** | **$\text{dBF}_{\text{pos}} = 50.05$** | **0.225 (0.112, 0.322)** | **$\text{dBF}_{\text{pos}} = 46.48$** | 0.019 (-0.012 to 0.048) | $\text{dBF}_{\text{pos}} = 9.696$ |
| | $k$ | -0.315 (-1.904, 0.648) | $\text{dBF}_{\text{neg}} = 1.769$ | -0.179 (-1.882, 0.898) | $\text{dBF}_{\text{neg}} = 1.378$ | -0.115 (-0.592, 0.265) | $\text{dBF}_{\text{neg}} = 2.827$ |
| Inverse Temperature | $y_0$ | **-16.180 (-18.521, -13.694)** | $\text{dBF}_{\text{neg}} = \text{Inf}$ | **-25.903 (-28.723, -23.249)** | $\text{dBF}_{\text{neg}} = \text{Inf}$ | **9.729 (7.518, 11.875)** | $\text{dBF}_{\text{pos}} = \text{Inf}$ |
| | $a$ | **-3.426 (-4.220, -2.598)** | **$\text{dBF}_{\text{neg}} = 49048.67$** | **-2.425 (-3.402, -1.490)** | **$\text{dBF}_{\text{neg}} = 7235.039$** | **-1.002 (-1.820, -0.235)** | **$\text{dBF}_{\text{neg}} = 107.328$** |
| | $k$ | -0.020 (-0.113, 0.070) | $\text{dBF}_{\text{neg}} = 2.083$ | **-0.173 (-0.283, -0.057)** | **$\text{dBF}_{\text{neg}} = 268.107$** | **0.153 (0.045, 0.255)** | **$\text{dBF}_{\text{pos}} = 150.137$** |

DV dependent variable, HDI highest density interval, $y_0$ intercept, $a$ asymptote, $k$ decay rate

